# Multicompartmentalized microvascularized tumor-on-a-chip to study tumor-stroma interactions and drug resistance in ovarian cancer

**DOI:** 10.1101/2024.05.29.596456

**Authors:** Simona Plesselova, Kristin Calar, Hailey Axemaker, Emma Sahly, Pilar de la Puente

## Abstract

**Introduction:** The majority of ovarian cancer (OC) patients receiving standard of care chemotherapy develop chemoresistance within 5 years. The tumor microenvironment (TME) is a dynamic and influential player in disease progression and therapeutic response. However, there is a lack of models that allow us to elucidate the compartmentalized nature of TME in a controllable, yet physiologically relevant manner and its critical role in modulating drug resistance.

**Methods:** We developed a 3D microvascularized multiniche tumor-on-a-chip formed by five chambers (central cancer chamber, flanked by two lateral stromal chambers and two external circulation chambers) to recapitulate OC-TME compartmentalization and study its influence on drug resistance. Stromal chambers included endothelial cells alone or cocultured with normal fibroblasts or cancer-associated fibroblasts (CAF).

**Results:** The tumor-on-a-chip recapitulated spatial TME compartmentalization including vessel-like structure, stromal-mediated extracellular matrix (ECM) remodeling, generation of oxygen gradients, and delayed drug diffusion/penetration from the circulation chamber towards the cancer chamber. The cancer chamber mimicked metastasis-like migration and increased drug resistance to carboplatin/paclitaxel treatment in the presence of CAF when compared to normal fibroblasts. CAF-mediated drug resistance was rescued by ECM targeted therapy. Critically, these results demonstrate that cellular crosstalk recreation and spatial organization through compartmentalization are essential to determining the effect of the compartmentalized OC-TME on drug resistance.

**Conclusions:** Our results present a functionally characterized microvascularized multiniche tumor-on-a-chip able to recapitulate TME compartmentalization influencing drug resistance. This technology holds the potential to guide the design of more effective and targeted therapeutic strategies to overcome chemoresistance in OC.

## Introduction

Ovarian cancer (OC) is a deadly disease representing the 5^th^ leading cause of cancer-related death among women in developed countries [1]. Due to the lack of specific symptoms in the early stage of the disease, it is mostly diagnosed in the advanced metastatic stage with a 5-year survival rate lower than 15% [1, 2]. Most patients are initially responsive to the standard-of-care taxane/platinum-based therapy, however, more than 80% of patients develop chemotherapy resistance within 5 years leading to elevated mortality and poor prognosis [1-3]. In the intricate landscape of OC biology, the tumor microenvironment (TME) has emerged as a dynamic and influential player in disease progression and therapeutic response [4, 5]. The TME encompasses a complex network of cellular and non-cellular components, including fibroblasts, imperfect vascularization, spatial heterogeneity in the distribution of nutrients, oxygen, and signaling molecules, and extracellular matrix (ECM), which collectively create a specialized niche supporting tumor growth. Recent advancements in OC research underscore the critical role of TME in modulating drug resistance, shedding light on the need to understand its compartmentalized nature [6-8]. The cellular crosstalk between tumor cells, stromal cancer-associated fibroblasts (CAF) and endothelial cells (EC) in the TME interferes with the cancer cell behavior, progression, and drug response [9, 10]. CAF are the major stromal cellular component surrounding the tumor core that interfere in pathogenesis, progression, invasion, angiogenesis, ECM remodeling, and drug resistance in many cancers [9, 10]. CAF actively participate in ECM remodeling by increased secretion of collagens through transforming growth factor-β (TGF-β) signaling converting the ECM into a stiff physical barrier for drug penetration and generating oxygen gradients that result in hypoxic niches [9, 11-14]. Interestingly, hypoxia, one of the key hallmarks of OC, is reciprocally implicated in enhanced TGF-β production and CAF activation, increasing the ECM remodeling [15, 16] and also enhancing tumor progression, metastasis, drug resistance, and angiogenesis [12, 17]. The imperfect tumor vasculature generated in the TME influences cancer progression, metastasis, pathogenesis, and it also aggravates hypoxia and causes decreased drug delivery into the tumor, thus having a negative impact in OC patient outcomes [10]. Therefore, a better understanding of the key components of TME compartmentalization and its profound implications for drug resistance in OC will ultimately pave the way for the development of more effective and personalized cancer therapies improving the survival rate in OC patients [18, 19].

Unfortunately, 95% of new cancer chemotherapeutics fail in clinical trials due to a lack of adequate preclinical models to mimic accurately the TME *ex vivo* [20-22]. Animal models remain a key asset in preclinical drug screening assays as they can recapitulate relevant TME components, but they are expensive, time-consuming, have associated ethical problems, and do not always represent the human TME and its key compartments and components [23, 24]. In that regard, recently the U.S. Food and Drug Administration (FDA) passed a law that no longer requires animal testing before human drug trials [25]. The traditionally used 2D cultures are easy to use, high-throughput, and inexpensive, although they are not able to recapitulate the intricacies of the TME, where factors like spatial heterogeneity, nutrient and oxygen gradients, and dynamic cellular interactions play pivotal roles in shaping drug responses [23, 24]. Three-dimensional (3D) cocultures in hydrogels can solve some of these limitations and conserve cellular crosstalk, and gradients but usually cannot recreate the compartmentalization and dynamics of the TME [23, 24, 26]. Moreover, spheroids and organoids are also a widely used model that can partially recapitulate the TME, but there is no fluid flow control or controlled compartmentalization, and the cells inside the spheroids are difficult to image, there is heterogeneity between samples, and not all cell types can naturally form them [23, 24, 26]. Microfluidic systems have emerged as a transformative tool, offering unparalleled insights into the dynamic interplay between tumors and therapeutic agents. Microfluidic systems or tumor-on-a-chip have unique capabilities to study drug resistance by recreating physiological conditions, including fluid shear stress, nutrient and oxygen gradients, and tissue spatial organization and cellular crosstalk, promoting a more accurate representation of the TME [24, 27].

Here, we have developed a multicompartmentalized microvascularized tumor-on-a-chip as a novel preclinical model that recapitulates key components of the dynamic and compartmentalized OC-TME to further evaluate their critical role in modulating OC drug resistance. We have validated our model for generation of oxygen gradients recapitulating the hypoxic OC-TME, formation of vessel-like structures recreating the TME vasculature and mimicking of aberrant ECM remodeling, processes that were enhanced by the presence of CAF in the stromal compartment. Moreover, we have performed drug response studies with standard-of-care treatment and addressed the CAF-induced drug resistance in OC by targeting the TGF-β signaling implicated in ECM remodeling. This enabling technology offers a more biomimetic environment for studying drug resistance offering unparalleled insights into the dynamics and complex compartmentalization of the TME.

### Material and Methods Microdevice fabrication

Compartmentalized tumor-on-a-chip was designed using KLayout 0.27.5 software with 5 chambers (2 mm x 4 mm) separated by trapezoidal posts with 100 µm gaps. The microdevice was manufactured using the SU-8 lithography technique by pouring polydimethylsiloxane (PDMS, Krayden Dow Sylgard 184 Silicone Elastomer Kit, NC9285739, FicherScientific) on SU8-2075 resin-covered wafer with UV-printed design and baking at 80°C for 2 hours. The height of the device was 100 µm. The inlets and outlets in the stromal and cancer chambers were cut with 1.5 mm punchers and in the circulation chambers with 3 mm Miltex® Biopsy punchers (Ted Pella, Inc). The PDMS design was bonded to a 24 x 60 mm coverslip (2-541-037, Fisher Scientific) using 115 V plasma cleaner (PDC-001, Harrick Plasma) at 0.5 Torr for 2 min. The devices were sterilized by UV light exposure for 30 min and baked overnight at 80°C to gain hydrophobicity.

### Cell culture

Ovarian cancer cell lines KURAMOCHI (RRID:CVCL_1345) and SKOV-3 (ATCC HTB-77), representing high-grade ovarian carcinoma and ovarian adenocarcinoma respectively, were cultured in specific media for ovarian cancer cells composed by DMEM (MT10013CV, Corning) and Ham’s F-12 (MT10080CV, Corning) media in 1:1 ratio supplemented by 10% (V/V) of fetal bovine serum (FBS, GibCo), 100 U/ml penicillin, and 100 mg/ml streptomycin (Corning CellGro). Human umbilical vein endothelial cells (HUVEC, ATCC CRL-1730) were cultured in F12K media (30-2004, ATCC) supplemented with 0.1 mg/ml heparin (H3393-100KU, Sigma Aldrich) and 30 µg/ml of endothelial cell growth supplement (ECGS, CB-40006, Fisher Scientific) for expansion and in VascuLife® VEGF media (1308000000000, Lifeline Cell Technology) for vessel formation. Primary human adipose-derived stem cells (StemPro^TM^, R7788115, ThermoFisher Scientific) were used as a normal fibroblasts control (NF) and they were cultured in MesenPro RS^TM^ Medium supplemented by MesenPro RS^TM^ Supplement (12746012, Gibco). Cancer-associated fibroblasts (CAF) specific for each OC cell line (CAF-KURAMOCHI or CAF-SKOV3) were developed by growing primary human uterine fibroblasts (HUF, ATCC PCS-460-010) in presence of conditioned media from each OC cell line as previously described [28].

### Multi-cultures in tumor-on-a-chip

Ovarian cancer cell lines were seeded into the central cancer chamber at a concentration of 4 x 10^6^ cells/ml in a 3D human plasma matrix based on fibrinogen crosslinking as we have previously described [29-31]. Briefly, to form the 3D plasma matrix, the cell suspension was mixed with human plasma from healthy donors, calcium chloride as a crosslinker reagent, and tranexamic acid as a stabilizer in 4:4:1:1 ratio, respectively. In the stromal chambers, 20 x 10^6^ cells/ml of EC were seeded in monoculture or coculture with 3 x 10^6^ cells/ml NF or CAF in a 3D human plasma matrix with 2 mg/ml Cultrex collagen I (3440-100-01, R&D Systems) in 1:1 ratio. In order to preserve the pressure differences inside the device, first, the stromal chambers were filled with cell suspension in the 3D matrix, and after crosslinking at 37°C for 10 min, the VascuLife® VEGF media was added in the circulation channel to allow vessel formation using 230 µl reservoir connectors (PG-RC-300UL-Q100, PreciGenome). The media was filled with a 2 cm difference (200 µl) between the inlets and the outlets of the circulation chambers to allow hydrostatic pressure and guarantee perfusion based on gravitational difference and interstitial and laminar flow of the fluid inside the microdevice [32]. Finally, the cancer chamber was filled with the OC cell suspension. As a control, a blank 3D matrix with no cells was injected into the stromal chamber. Cell-specific media was added in the inlets and outlets of each chamber to ensure proper media and nutrient gradients across the chip.

### Hypoxia studies

KURAMOCHI cells (3x10^6^ cells/ml) were cultured for 7 days in the central chamber with a blank 3D matrix, EC in monoculture, or EC in coculture with NF or CAF in the stromal chamber as described above. Image-iT^TM^ Green Hypoxia Reagent (5 μM, EX 488/ EM520 nm, I14833, Invitrogen) was added for 24 hours in the circulation chamber to detect the cells with oxygen levels below 5% and then washed with PBS 1x for 8 h. The images were taken using Nikon A1R Ti2E confocal microscope (EX 488 nm, EM 525 nm) with 10x magnification using large image acquire mode with stitching with 10% overlapping through optimal pathway at 1024x1024 pixel resolution.

### Collagen I staining

KURAMOCHI cells (3x10^6^ cells/ml) were cultured for 7 days in the presence of a blank scaffold, and monoculture EC, NF, or CAF in the stromal chamber. Microchips were washed with PBS 1x, fixed with 1% paraformaldehyde, washed again with PBS 1x, and blocked with 4% (w/V) BSA in VascuLife® VEGF media. Finally, the tumor-on-a-chip was incubated with AF488-anti-collagen I antibody (1:1000, ab275996, Abcam) and DAPI 1 µg/ml (62248, ThermoScientific^TM^) in VascuLife® VEGF media supplemented with 0.1% (w/V) BSA (A2508, Sigma-Aldrich) for 24 h, next washed with PBS 1x and imaged on Nikon A1R Ti2E confocal microscope (EX 488 nm, EM 525 nm). Close-up images were taken in the different compartments of the device. Then, the 3D scaffolds inside the microfluidic device were digested with 20 mg/ml collagenase I (Gibco^TM^, USA) for 3 h, and imaging flow cytometry was performed on Cytek^®^ Amnis^®^ ImageStream®^X^ MkII Imaging Flow Cytometer (Cytek Biosciences) using the INSPIRE software at 40x magnification at low speed. Images were acquired with a 488 nm laser power of 35 mW, a 785 nm laser at 2 mW for side scatter and in brightfield.

### Drug penetration and uptake

For drug penetration assays, the tumor-on-a-chip was filled with a blank 3D matrix and incubated for 30 min at 37°C for crosslinking. For drug uptake assays the microchips were filled with SKOV-3 cells (3x10^6^ cells/ml) in the cancer chamber surrounded by a blank 3D scaffold, EC, or EC in coculture with NF or with CAF in the stromal chamber as described above. The cells were grown for 5 days to allow ECM remodeling and tumor-stroma interactions. For both drug penetration and drug uptake assays, doxorubicin 100 µM (100280, MedKoo Biosciences), a naturally fluorescent drug, was added in both circulation chambers using the reservoir connectors. The images were taken at different time points (0; 0.25; 0.5; 0.75, 1; 1.25; 1.5; 1.75; 2, 2.5; 3; 4; 5; 6; 7; 8; 24 hours) for drug penetration and at 24 h for drug uptake using Nikon Eclipse Ni-E fluorescent microscope (EX 559.5 nm/EM 645 nm) at 10x magnification.

### Migration studies

OC cell lines (KURAMOCHI and SKOV-3) were stained with surface membrane marker Invitrogen^TM^ DiD (D307, Invitrogen) and seeded at 4x10^6^ cells/ml concentration in the cancer chamber surrounded by blank 3D scaffold, endothelial cells stained with BV605-anti-CD31 antibody (303122, Biolegend) in monoculture or co-culture with either NF or CAF detected with FITC-anti-CD90 antibody (328108, Biolegend) in the stromal chamber. VascuLife® VEGF media was changed daily to allow vessel formation. The microfluidic devices were imaged on days 3, 7, and 14 using fluorescent confocal microscope Nikon A1R-Ti2E at 10x magnification with 1024x1024 pixel image resolution and in large image acquiring mode with 10% overlapping through the optimal pathway.

### Drug treatment

KURAMOCHI or SKOV-3 cells (4x10^6^ cells/ml) were seeded in the cancer chamber in the presence of a blank 3D scaffold, EC (20x 10^6^ cells/ml) or EC in coculture with NF or CAF in the stromal chamber for 3 days to allow cellular crosstalk and ECM remodeling. Then, the standard-of-care chemotherapeutic drugs paclitaxel (12.5 nM, S1150, Selleckchem) and carboplatin (25 µM, 100130, MedKoo) were added into both circulation chambers for 4 days. For ECM remodeling targeting studies, the cells were pre-incubated with halofuginone 10 nM (S8144, SelleckChem), a TGF-β signaling inhibitor for 72 h, and then paclitaxel/carboplatin/halofuginone combo was added for 4 days. A microfluidic device treated with DMSO was used as a control for cell viability. The day before analysis, the cells were stained with Nuclear ID^®^ Blue/Red cell viability kit (ENZ-53005-C100, Enzo Life Sciences) and images were taken in Nikon Eclipse Ni-E fluorescent microscope at 10x magnification in DAPI (EX 383-408 nm/EM 435-485 nm) and TexRed channels (EX 540-580 nm/EM 593-668 nm).

### Image data analysis

Fluorescent images acquired on Nikon Eclipse Ni-E or confocal microscope Nikon A1R-Ti2E, unless otherwise stated, were processed, and analyzed with NIS-Elements AR Analysis 5.21.03 Software. Binary thresholding was used to detect the cells of interest. For the vessel’s characterization, area restriction in a range of 50% higher than the calculated mean value of all objects and the largest object area value (3,000 µm^2^-50,000 µm^2^) was applied to detect vessel-like structures of EC in TexRed channel (BV605-anti-CD31 antibody). Circularity (0-1), elongation, and area (μm^2^) parameters were determined by automatic measurement of selected objects and the inner diameter of the vessels (μm) was manually determined with the distance measurement tool. In the collagen I secretion studies, the unspecific green background from the 3D scaffold was subtracted in the fluorescent images of AF488-anti-collagen I, and the MFI of AF488 inside individual cells was measured with automatic measurement. The images taken on ImageStream®^X^ MkII Imaging Flow Cytometer were analyzed using IDEAS v6.2 analysis software. Each sample was processed by first using the area of the brightfield and the brightfield aspect ratio to identify single round cells discarding debris and speed beads. Then, the brightfield gradient RMS value was used to gate in-focus events for further analysis. Finally, events with no Mean Pixel AF488 intensity were excluded from the analysis discarding debris (Supplementary Fig. 1a). The area of each cell was calculated using a brightfield-based object mask and the AF488-colagen I intensity was calculated from the pixel intensity within the cell mask (Supplementary Fig. 1b). All images show the median AF488-collagen I intensity for each sample and were window-leveled based on the median intensity of the CAF images. Finally, the MFI of AF488-collagen I was analyzed in individual cells in FlowJo_v10.8.1_CL software and representative histograms were generated for each condition. In the oxygen gradients studies, the tumor-on-a-chip was divided into 9 equal vertical portions from left to right and the mean fluorescent intensity of the Image-iT^TM^ green hypoxia reagent was measured inside individual cells through binary thresholding and automatic detection of MFI. In the migration studies, binary thresholding was used to detect the KURAMOCHI cells, and the number and area of the clusters were measured in the different conditions using automated measurement and applying specific restrictions of area in a range of 50% higher than the calculated mean value of all objects in cancer chamber and the largest object area value (1,300 µm^2^ – 90,000 µm^2^) (supplementary Fig. 2). To characterize the metastasis-like migration of SKOV-3 cells towards the stromal chamber, the migration distance was measured manually using the distance measurement tool and MFI of DiD-marked OC cells (Cy5 channel) was determined by automatic measurement feature in rectangular region of interest (ROI) selected in the stromal chamber of the device to detect the number of cells that migrated. For drug penetration studies, the MFI of doxorubicin was quantified in stromal and cancer chambers separately using rectangular ROI selection and for drug uptake images, the red background of doxorubicin penetration into the 3D matrix was subtracted to detect intracellular drug fluorescence. The representative pictures in the drug treatment studies were processed adjusting the blue staining for live cells into green color for better visualization and the number of live (green) and dead (red) cells was determined by manual counting in ImageJ software using the cell counter plugin.

### Statistical analysis

Experiments were performed in triplicates and repeated at least three times. All graphs were generated and analyzed using GraphPad Prism 9 software as Mean ± SD, and statistical significance was analyzed using t-test, One-way or Two-way ANOVA, and p-value lower than 0.05 was considered significant.

## Results

### Multiniche microvascularized tumor-on-a-chip recapitulates the compartmentalized OC-TME

To develop a new physiologically relevant model to mimic the spatial organization and the biological compartments of the OC-TME, we have designed a novel microvascularized multicompartmentalized PDMS tumor-on-a-chip. Specifically, this device consisted of a central chamber to seed OC cell lines, flanked by two stromal chambers on both sides to grow EC in monoculture or coculture with NF or CAF to study tumor-stroma crosstalk and the influence of TME compartmentalization in drug resistance in OC. Two outer circulation channels on each side were included to perfuse media or drugs into the device. The dimensions of each chamber were 2 mm x 4 mm x 0.1 mm. All compartments were separated by trapezoidal pillars with 100 µm gaps to maintain the compartmentalization, and allow cell-cell interactions, migration, and media and drug flow (Fig. 1a). Tumor-on-a-chip was functionally characterized for the recreation of key physical components of the complex OC-TME. First, the design was measured for the formation of vascularization in the stromal chambers surrounding the cancer chamber. Vessel-like structures were formed in the three experimental groups including EC, either in monoculture or coculture with NF or CAF. No significant differences were observed in the circularity and area of the vessels between the groups, although the presence of CAF significantly increased the inner diameter of the vessels from 60 µm to 80 µm and the elongation from 1.9 to 2.1 when compared to EC monoculture (Fig. 1b). Second, the tumor-on-a-chip was evaluated for the recapitulation of ECM remodeling associated with tumor growth and driven by stromal cells, which is a key contributor to reduction of therapeutic efficacy. Collagen I secretion, as the most common and abundant ECM protein in the OC-TME, was assessed by immunofluorescence staining both by confocal imaging and imaging flow cytometry providing high throughput single cell measurements coupled with single cell imaging. While confocal images revealed no collagen I expression in blank, OC and EC cells, monoculture NF and CAF had positive collagen staining and CAF had significantly higher expression measured by MFI than the NF (Fig. 1c). Confocal imaging of whole chips had significant matrix background when stained for collagen, therefore, cells were isolated from the device and analyzed by imaging flow cytometry. Similarly, collagen I expression was significantly higher in CAF compared to OC, EC, and NF (Fig. 1d). Finally, we evaluated the tumor-on-a-chip for the recapitulation of key spatial tissue gradients, particularly for oxygen gradients allowing to recreate the hypoxic OC-TME. The four experimental groups were stained with green hypoxia ImageiT^TM^ reagent that fluoresces when cells are exposed to less than 5% oxygen. A hypoxic niche core in the central cancer chamber was validated in all 4 conditions, but only the incorporation of CAF in the stromal chamber allowed for a spatial tissue oxygen gradient from the circulation chambers towards the cancer chamber (Fig. 1e). These findings validated our model as suitable for recreating the compartmentalized OC-TME and the influence of CAF in vessel-like structure formation, ECM remodeling, and spatial oxygen gradients.

**Fig. 1.**
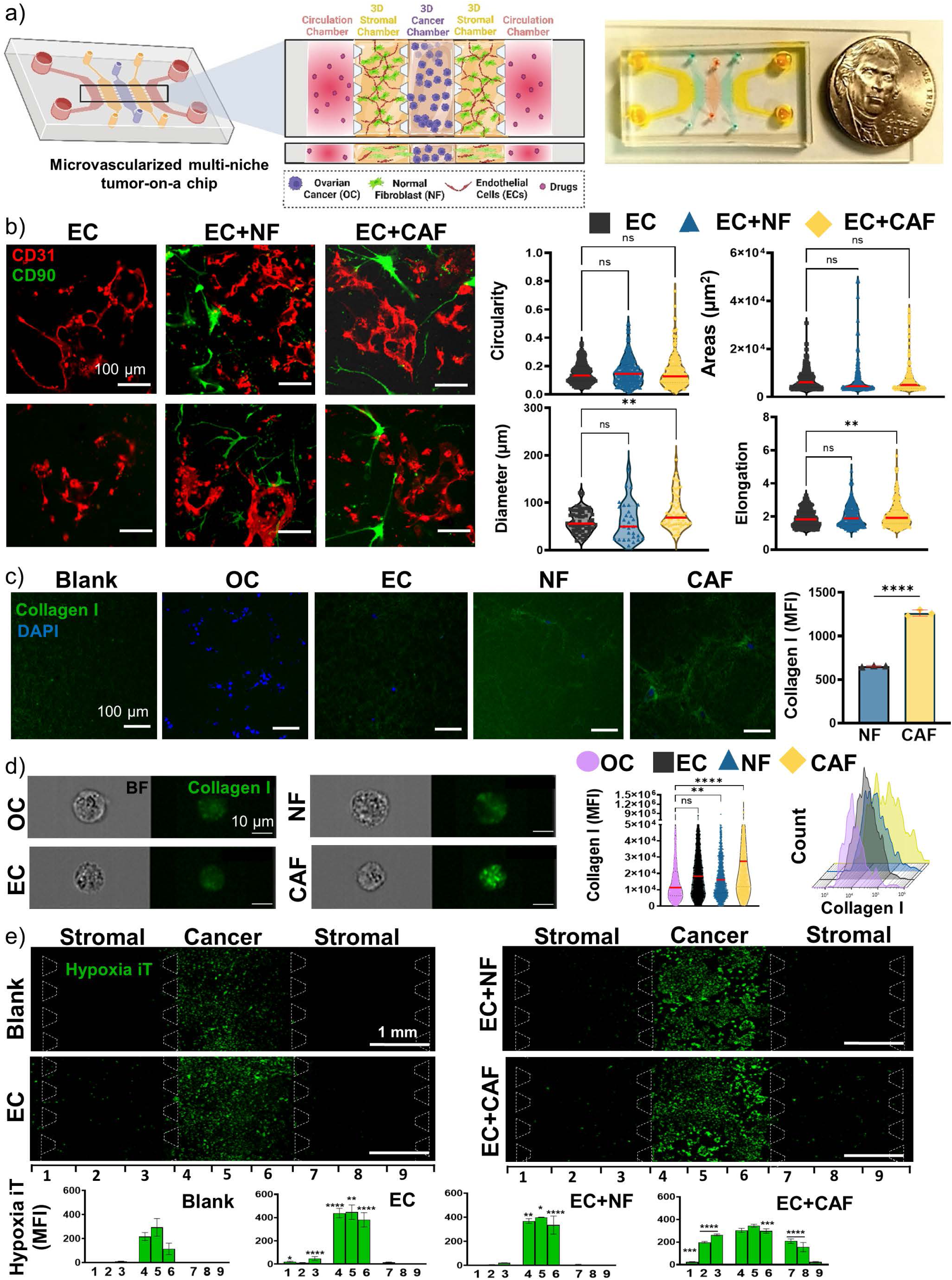
Multiniche microvascularized tumor-on-a-chip recapitulates the compartmentalized OC-TME. a) Schematic representation of tumor-on-a-chip device and close-up view of the different chambers and cell types seeded inside (left) and real image of microfluidic device filled with dyes next to a nickel representing the size of the platform (right). **b)** Representative images of the stromal chamber with vessel-like structures formation of endothelial cells (EC) stained with BV605-anti-CD31 (red) in monoculture or coculture with normal fibroblasts (NF) or cancer-associated fibroblasts (CAF) stained with FITC-anti-CD90 (green) and the quantification of the circularity, area, inner diameter, and elongation of the vessels. Median value is marked in red. Scale Bar=100µm. *p<0.05, One-Way ANOVA compared to EC. **c)** Representative images of the stromal chamber by immunofluorescent staining with AF488-anti-collagen I in blank gel or mono-culture cells grown in the microfluidic device for 7 days and quantification of the mean fluorescent intensity (MFI) of AF488. Scale Bar= 100µm. Mean±SD, **** p<0.0001, t-test. **d)** Imaging flow cytometry analysis of cells isolated from microfluidic device and analyzed for collagen I, including representative images, representative histogram of each condition and violin plot for quantification of MFI of AF488 inside individual cells. Median value is marked in red. **p<0.01, ****p<0.0001, Two-Way ANOVA compared to blank. Scale Bar=10µm. **e)** Representative images of OC cells grown in presence of blank scaffold, EC, EC+NF or EC+CAF in the stromal chambers for 7 days and stained with hypoxic Image-iT^TM^ reagent (green) and quantification of MFI of the device divided in 9 sections from left to right. Scale Bar=1mm. Mean±SD, *p<0.05, **p<0.01, ***p<0.001, **** p<0.0001, Two-Way ANOVA compared to Blank.

### Tumor-on-a-chip allows for cluster formation and metastasis-like migration while recapitulating tumor-stroma interactions

To study cancer cell behavior and cellular crosstalk, tumor-on-a-chip devices were monitored over 14 days in culture. On one hand, KURAMOCHI, a non-metastatic high-grade serous OC cell line that forms clusters in 3D culture [33] was evaluated and found that over time KURAMOCHI formed more and bigger clusters and established cell interactions with the stromal compartment (Fig. 2a). Numbers of cluster and cluster size were significantly higher in OC cells when no stroma was present (blank) on days 3 and 7. In the presence of stromal cells on day 3, the number and area of clusters surrounded by CAF were higher than in EC and NF. While day 7 showed a significantly elevated formation of clusters in the presence of CAF and EC compared to NF, day 14 had a similar number of clusters in all conditions even though NF and CAF had significantly higher areas compared to blank (Fig. 2b). On the other hand, SKOV-3, a serous adenocarcinoma very aggressive and metastasis-like cell line [34, 35], was also monitored and identified that formed tumor-stromal interactions and initiated metastasis-like migration towards stromal chambers on days 7 and 14 (Fig. 2c). The distance of migration of the OC cells was measured and while there were no significant differences on day 7, migration was significantly reduced on day 14 in the conditions that contain stromal fibroblasts (NF and CAF) compared to blank. When no stroma was included, the SKOV-3 cells migrated longer distances than when stroma cells were present (335±33 µm on day 7 and 663±54 µm on day 14), followed by SKOV-3 cultured with EC (280±24 µm on day 7 and 349±26 µm on day 14). When fibroblasts were seeded in the stromal chambers, CAF allowed higher migration of SKOV-3 cells compared to NF (216±21 µm on day 7 and 273±33 µm on day 14 compared to 188±23 µm on day 7 and 258±32 µm on day 14 in NF) (Fig. 2d). Also, the number of SKOV-3 cells that migrated towards stromal chambers determined by MFI measurement of DiD-stained cells in the stromal compartment was higher when no stroma was present, and EC and CAF induced significantly more metastasis-like migration than NF.

**Fig. 2.**
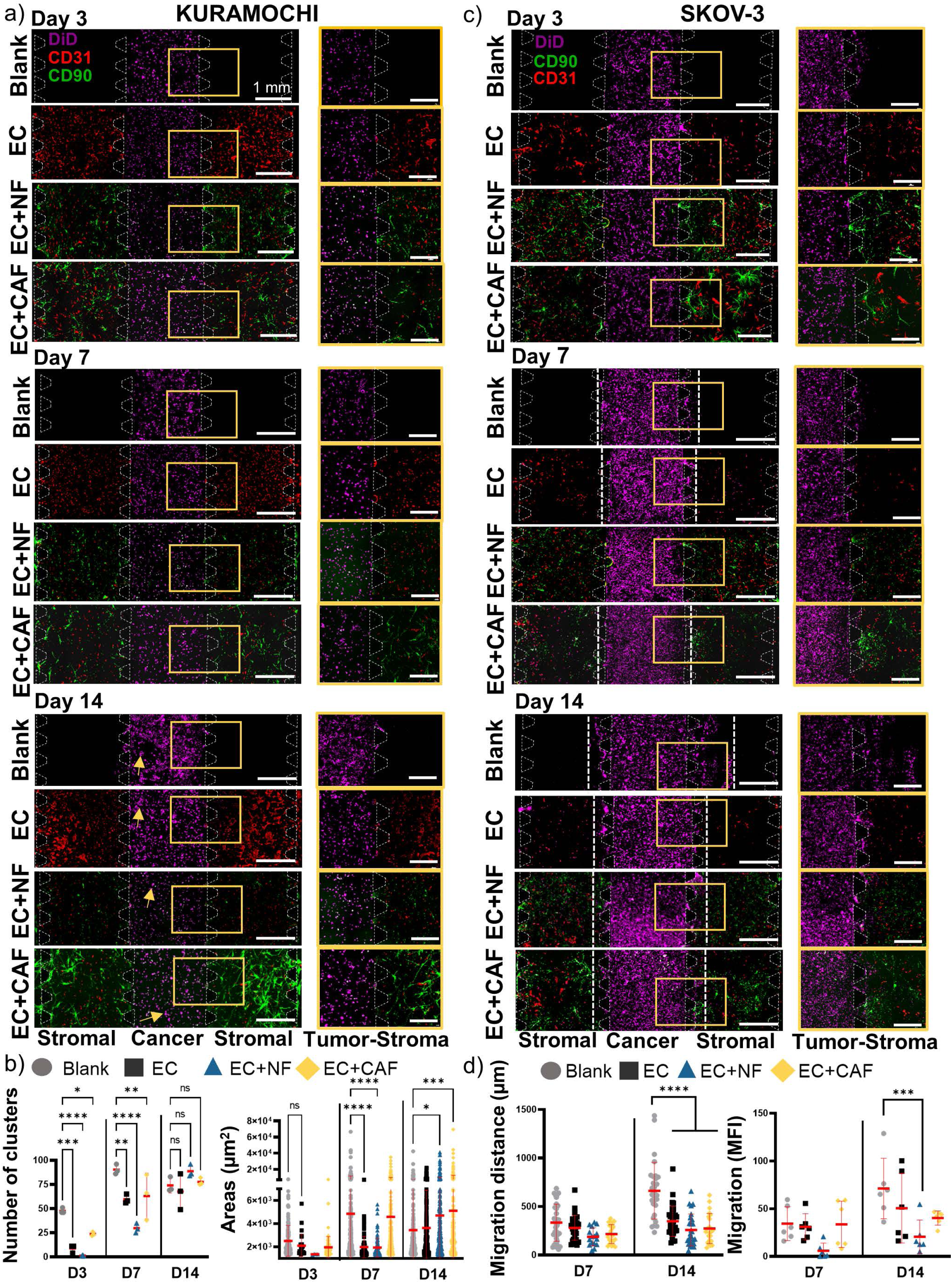
Tumor-on-a-chip allows for cluster formation and metastasis-like migration while recapitulating tumor-stroma interactions. a) KURAMOCHI cells (DiD, purple) were grown in cancer chamber with blank scaffold, EC (BV605-anti-CD31, red), EC+NF (FITC-anti-CD90, green) or EC+CAF (FITC-anti-CD90, green) in stromal chambers and images were taken on days 3, 7 and 14. Scale bar=1 mm. Close-up magnification images show the interactions between cancer and stromal chambers (yellow rectangles) and yellow arrows show the cluster formation on day 14. Scale Bar=0.5mm. **b)** Quantification of number of clusters and cluster area for KURAMOCHI cultures (low-metastatic potential). Mean±SD, *p<0.05, **p<0.01, ***p<0.001, **** p<0.0001, Two-Way ANOVA compared to blank. **c)** SKOV-3 cells (DiD, purple) were grown in cancer chamber with blank scaffold, EC (BV605-anti-CD31, red), EC+NF (FITC-anti-CD90, green) or EC+CAF (FITC-anti-CD90, green) in stromal chambers and images were taken on days 3, 7 and 14. Scale bar=1 mm. Close-up magnification images show the interactions between cancer and stromal chambers (yellow rectangles). Scale bar=0.5 mm. White discontinued lines denote distance of migration towards stromal chamber. **d)** Quantification of distance of migration and number of migrated cells (MFI of DiD) for SKOV-3 cultures (high metastatic potential). ***p<0.001, **** p<0.0001, Two-Way ANOVA compared to blank.

### Microfluidic device mimics drug penetration gradients in OC-TME

To analyze the fluid flow inside of our tumor-on-a-chip, a blank 3D scaffold (without cells) was injected into cancer and stromal chambers, and once polymerized, doxorubicin was injected in both circulation channels and fluorescent time-lapse microscopy was performed (Fig. 3a). A laminar flow of doxorubicin (red) can be seen on the representative images in Fig. 3b over time, reaching full penetration of the whole central chamber at 24 h. The doxorubicin penetration was quantified by measuring its MFI at different time points. As expected, doxorubicin penetrated faster in the stromal chambers at around 2-4 h, but there was a delay in drug penetration into the central chamber (7 h), recapitulating tumor drug penetration. Importantly, both stromal and cancer chambers reached plateau full drug penetration at 24 h (Fig. 3c). Having this in mind, SKOV-3 cells were seeded in the central chamber in the presence of blank scaffold, EC, or EC in coculture with NF or CAF and grown for 5 days to allow TME spatial compartmentalization including vessel formation, ECM remodeling and oxygen gradients, and then doxorubicin was injected in both circulation channels and fluorescent images were taken at 24 h (Fig. 3d). As observed in Fig. 3e and in the supplementary video, complete drug penetration and uptake in cells were reached at 24 h, a time point that was established as a minimal time for drug screening incubations.

**Fig. 3.**
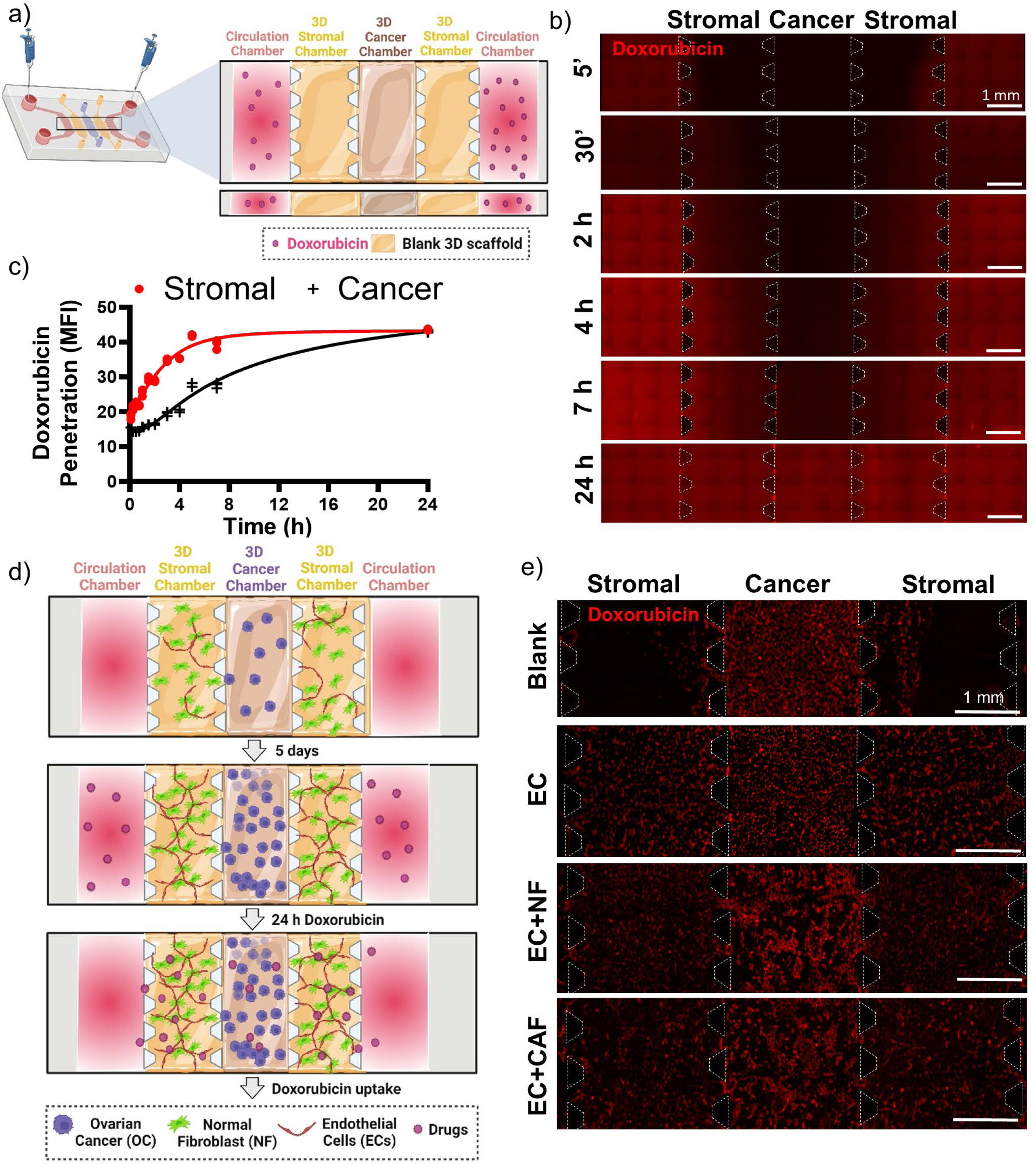
Microfluidic device mimics drug penetration gradients in OC-TME. a) Schematic representation of method used for doxorubicin penetration assay, where doxorubicin (100 µM) was injected in both circulation channels and images were taken in time-lapse fluorescent microscope up to 24 h. **b)** Representative images at 5 min, 30 min, 2, 4, 7 and 24 h of doxorubicin penetration. Scale Bar=1mm. **c)** Quantification of MFI of doxorubicin penetration inside the cancer and stromal chambers at each time point. **d)** Schematic representation of method used for doxorubicin uptake, cancer cells were cultured surrounded by different stromal cells (EC, NF, or CAF) or blank scaffold as a control for 5 days and then doxorubicin (100 µM) was injected in both circulation channels and images were taken at 24 h. **e)** Representative images of doxorubicin uptake by cells at 24 h. Scale Bar=1mm.

### Multiniche microvascular tumor-on-a-chip recapitulates CAF-induced drug resistance in OC-TME

The microfluidic model was evaluated for drug screening and to study the influence of TME compartmentalization in drug resistance in OC. OC cell lines were grown in the absence of stroma (blank 3D scaffold) or presence of the different stromal cells (EC, EC in coculture with NF or CAF) for 3 days to allow spatial TME compartmentalization (vascularization, oxygen gradients and ECM remodeling), then, the devices were treated with a combination of standard-of-care chemotherapeutic drugs paclitaxel and carboplatin (PC) (Fig. 4 a). Representative close-up images of the central chamber seeded with KURAMOCHI or SKOV-3 cells are shown in Fig. 4b and Fig. 4d, respectively. There were no significant dead cells observed in either of the stromal cells as seen in Supplementary Fig 3. The number of live (green) and dead (red) cells was determined, and drug cytotoxicity for KURAMOCHI (Fig. 4c) and for SKOV-3 cells (Fig. 4e) was expressed as a fold change of the percentage of dead cells compared to DMSO control treated condition. In both cell lines, in the absence of stroma (blank) or presence of normal stroma (EC or EC+NF), there was a significant cytotoxic effect of the paclitaxel/carboplatin combination, but in the presence of CAF, there was no killing effect observed compared to DMSO treated condition, validating our model to recapitulate the influence of stromal cells in drug response in OC cancer, specifically CAF-induced drug resistance.

**Fig. 4.**
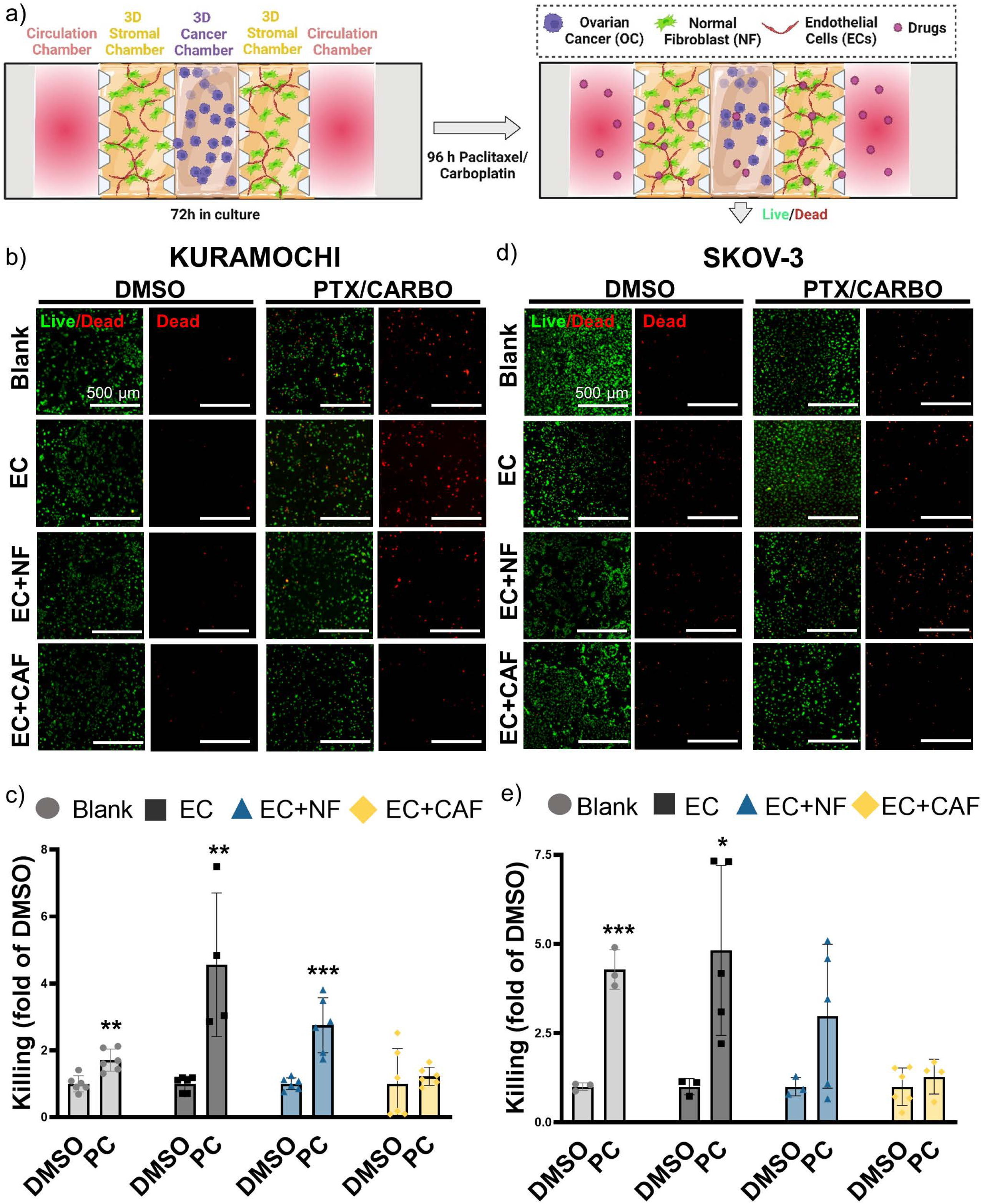
Multiniche microvascular tumor-on-a-chip recapitulates CAF-induced drug resistance in OC-TME. a) Schematic representation of the methods used in this experiment where OC cell lines were cultured in presence of different stromal components (blank 3D scaffold, EC, EC+NF and EC+CAF) in stromal chamber for 3 days to allow compartmentalization and then treated with paclitaxel and carboplatin (PC) for 4 days. DMSO-treated chips were used as a control. Nuclear ID Live (green)/dead (red) reagent was used to determine cell viability. **b)** Representative images of merged green(live)/red(dead) and red channel (dead cells) for KURAMOCHI in cancer chamber. Scale Bar=500µm. **c)** Quantification of fold change of percentage of cell death in DMSO treated condition. Mean±SD, **p<0.01, *** p<0.001, t test to corresponding DMSO control. **d)** Representative images of merged green(live)/red(dead) and red channel (dead cells) for SKOV3 in cancer chamber. Scale Bar=500µm. **e)** Quantification of fold change of percentage of cell death of DMSO-treated condition. Mean±SD, *p<0.05, *** p<0.001, t test to corresponding DMSO control.

### CAF-mediated drug resistance can be rescued by an anti-TGFb targeting agent

We used halofuginone as a targeting agent inhibiting TGF-β signaling to overcome the drug resistance in OC caused by CAF. KURAMOCHI cells were seeded in the central chamber surrounded by a blank scaffold, EC, EC in coculture with NF or CAF in two sets of microdevices. One set of the microfluidic devices was grown untreated to allow spatial TME compartmentalization and the other was pre-incubated with halofuginone in order to avoid TGF-β-mediated ECM remodeling by CAF. After 3 days, paclitaxel/carboplatin combination was added to the first set and paclitaxel/carboplatin/halofuginone to the second set of microfluidic devices for another 4 days. Then, cells were stained with viability reagent as previously described (Fig. 5a). Representative images of the central chamber of both sets, paclitaxel/carboplatin-treated (left) and paclitaxel/carboplatin/halofuginone-treated (right) are shown in Fig. 5b and revealed increased death in the co-culture CAF condition with halofuginone when compared to standard of care alone treatment. No significant dead cells were observed in the stromal chambers as seen in the representative images of whole microfluidic devices in Supplementary Fig 4. The cytotoxic effect was expressed as a fold change of paclitaxel/carboplatin-treated condition and as seen in Fig. 5c where halofuginone did not have any effect on the cancer cells grown in the presence of normal stroma, however, it rescued the CAF-mediated drug resistance to standard of care drugs.

**Fig. 5.**
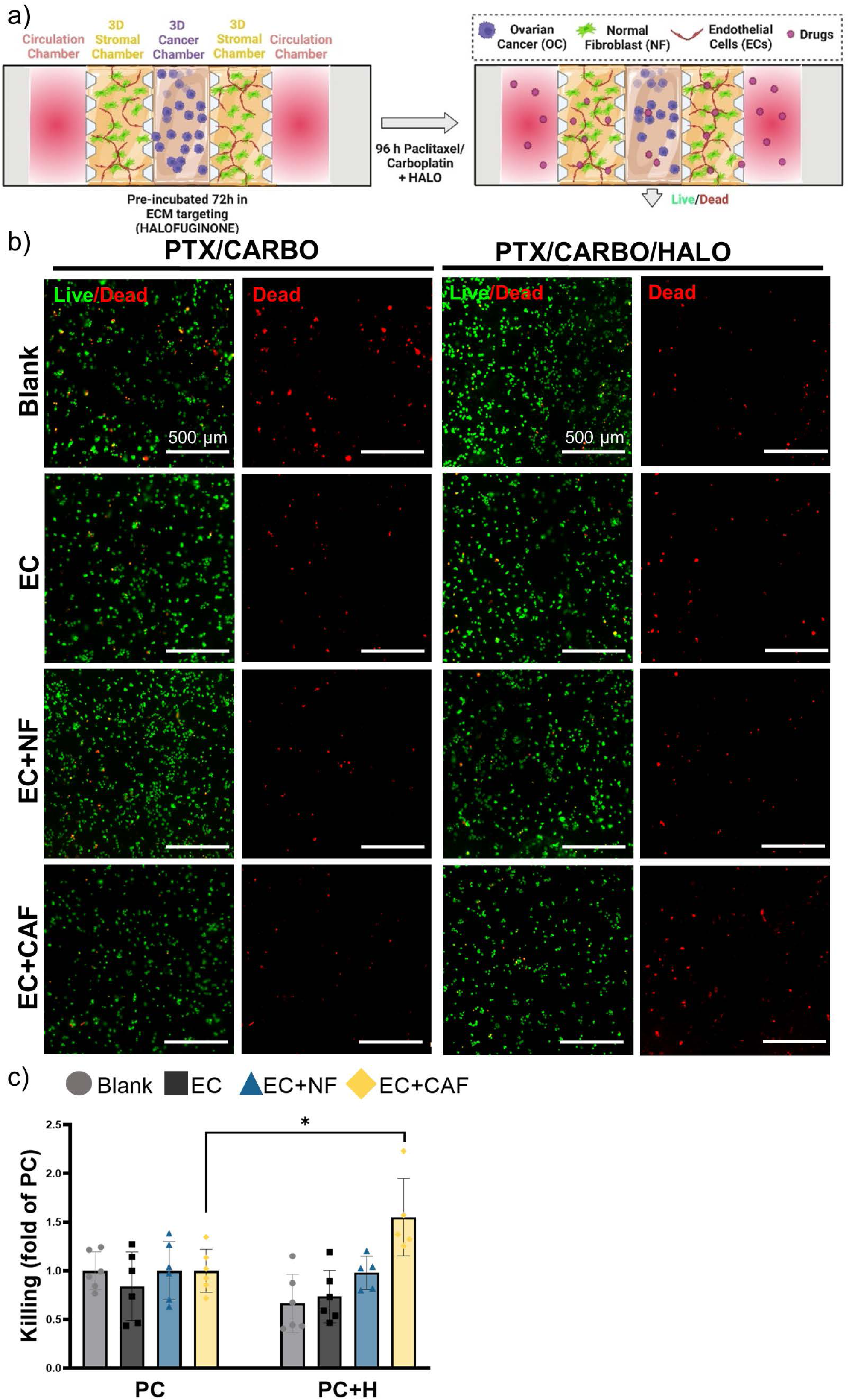
CAF-mediated drug resistance can be rescued by an anti-TGFb targeting agent. a) Schematic representation of methods used in this approach where the chips were pre-treated with halofuginone (HALO) as a targeting agent for TGF-β then treated with paclitaxel and carboplatin (PC) for 4 days**. b)** Representative close-up images of KURAMOCHI grown in central chamber with blank scaffold, endothelial cells (EC), EC in co-culture with NF or CAF in stromal chamber treated with PC or PC with halofuginone (PC+H) (as pre-treatment). Live/Dead Nuclear ID dye was used to detect live cells (green) and dead cells (red). Scale Bar=500µm. c) Quantification of fold change of percentage of cell death of PC condition. Mean±SD, *p<0.05, t test to corresponding PC condition.

## Discussion

In this study, we have developed a versatile multi-compartmentalized microvascularized PDMS tumor-on-a-chip, where spatial TME compartmentalization was recapitulated including vascularization, ECM remodeling, oxygen gradients, as well as drug penetration and uptake kinetics into tumor. Importantly, the device allowed us to investigate tumor-stroma cellular crosstalk, compare normal or cancer-associated fibroblast’s role in modulating drug response, and further investigate targeted therapies that can overcome this induced drug resistance. This study demonstrates that CAF are key contributors in the OC-TME compartmentalization and influence drug response, which is in accordance with ours and other reports [28, 36-38]. This device grants drug screening under a faithful recapitulation of the OC-TME, which would provide strong evidence for allowing new opportunities to improve chemotherapy effectiveness and mitigating chemoresistance in ovarian cancer.

Ovarian tumors are heterogeneous where the cancer-stroma interplay is important in tumor progression and therapeutic response and it is vital to mimic the cellular and acellular components of the TME, and its spatial organization in preclinical models in order to find new therapies and improve the outcomes of OC patients [39]. In terms of key cellular compartment recapitulation for mimicking the OC-TME compartmentalization, this study has used two distinct and valuable OC cell lines. We used the KURAMOCHI cell line, as a model for high-grade serous ovarian carcinoma (HGSOC) which is the most common and deadliest subtype of OC [1, 33], and the SKOV-3 cell line as a model for serous ovarian adenocarcinoma, as the most used cell line in OC preclinical research [34, 35]. To model the stromal component of the OC-TME, we used endothelial cells, or their co-culture with normal fibroblasts or cancer-associated fibroblasts. Specifically, we used human umbilical vein endothelial cells (HUVEC) which are widely used in vascularization microfluidic studies because of their easy growth in 3D scaffolds and formation of vessel-like structures [40-45], even though they do not represent exactly the behavior of the native EC in OC tumors, which could be included in future studies [46]. Moreover, human adipose-derived stem cells (ADSC) were used as a normal fibroblast model because of their high expansion ability and reproducibility suitable for high-throughput research, although they are not directly representative of normal tissue origin including fallopian tube or ovary [47]. Currently, the OC-stroma interaction studies use a variety of normal fibroblasts from different non-ovarian origins such are NIH3T3 fibroblasts from mouse embryos [28], lung fibroblasts [48], or dermal fibroblasts [49]. Unfortunately, the immortalized human normal ovarian fibroblast cell line (NOF151-hTERT) [50] is not commercially available. Therefore, we selected ADSC as it is demonstrated that human mesenchymal stem cells are functionally resting normal fibroblasts having similar phenotypes and characteristics and are widely used in bioengineering studies [51, 52]. Moreover, it is known that in the TME cancer cells recruit stromal cells into their vicinity through tumor-stromal interactions and signaling molecules and activate them into cancer-like phenotype [36]. In this study, we have used cancer-associated fibroblasts, as activated fibroblasts, derived from primary human uterine fibroblasts by their exposure to conditioned media from specific OC cell line (KURAMOCHI or SKOV-3) that have been previously functionally characterized in terms of CAF-like phenotype, higher proliferation rate, contractility, ECM remodeling, tumor progression, and drug resistance [28]. CAF are the most important cell type in the OC-TME constituting a heterogeneous population of cells because they can be derived from normal fibroblasts, bone marrow mesenchymal stem cells, adipocytes, endothelial cells, or epithelial cells and they can be activated through different signaling pathways, exosomes, or cytokines such as TGF-β and also through ECM remodeling, although most of these CAF subtypes have not been fully characterized yet. Therefore, using a well-characterized cell source is important to avoid the heterogeneity from primary CAF cultures that also have low reproducibility, low growth rate, and finite life span, thus they are not suitable for high throughput studies.

Once CAF in the TME are activated, they interact with the cancer cells, other stromal cells, and ECM through different signaling molecules, cytokines, chemokines, and exosomes, and participate in tumor progression, invasion, metastasis, angiogenesis, ECM remodeling and drug resistance [26, 37, 53, 54]. Even though the OC drug resistance mechanisms remain still unclear, it is known that CAF may play an important role as they are the key component of the OC stroma implicated in ECM remodeling [9]. Activated CAF secrete ECM proteins such are collagens mediated by TGF-β signaling, converting the ECM into a stiff obstacle for drug penetration and generating gradients of oxygen that result in a hypoxic tumor core [36, 37]. Hypoxia also enhances drug resistance by inducing different changes in gene and protein expression affecting many cellular and physiological functions, resulting in poor prognosis of OC patients [6, 12, 37, 54]. CAF-induced ECM remodeling can compress the vessels in the TME reducing drug delivery inside the tumor and CAFs also secrete hypoxia-induced angiogenesis regulator (HIAR) and vascular endothelial growth factor (VEGF-A) promoting aberrant and exacerbated angiogenesis [54]. Furthermore, CAF enhance vascular permeability and leakage that impede drug penetration into the tumor by different factors such are VEGF, lipoma preferred partner (LPP), and platelet-derived growth factor receptor (PDGFR) [54]. CAF also interfere with drug resistance by inducing and maintaining the stemness of the tumor, promoting increased efflux of chemotherapeutic drugs, and by secreting pro-inflammatory cytokine IL-6 that has been correlated with resistance to paclitaxel [8, 54, 55]. CAF are also implicated in the induction of epithelial-mesenchymal transition (EMT) increasing the invasiveness of the cancer cells and cisplatin resistance [56]. Moreover, CAF can reprogram metabolically the cancer cells to enhance their survival, and they secrete long non-coding RNA ANRIL that suppresses the expression of drug transporters to promote drug efflux in OC and induce cisplatin resistance [54, 56, 57]. Stromal cells from ascites have been reported to confer drug resistance to OC cells, as the cancer cells acquire the functional p-glycoproteins for drug efflux through trogocytosis of the stromal cell’s membrane [58]. Current therapeutic strategies to overcome drug resistance induced by CAF consist in targeting (I) the cells of origin of CAF, (II) CAF markers (FAP), (III) CAF activation signaling pathways, (IV) or CAF-directed therapeutic delivery [54]. Also, the treatment with chemotherapeutics in OC can stimulate TGF-β secretion and induce more drug resistance [59], therefore targeting TGF-β signaling could improve the OC patient outcomes. For example, halofuginone, an anticoccidial drug, is a TGF-β signaling inhibitor that had antitumor effects and reduced collagen synthesis in animal models of several solid cancers (lung, melanoma, breast, pancreas, etc.) [60], and had synergetic effect with chemotherapeutic drugs overcoming chemotherapeutic resistance in lung, prostate, and colorectal cancers, and it is currently explored in several clinical trials in bladder cancer, AIDS-related Kaposi sarcoma, and in advanced solid tumors [61]. Geyer *et al*. used halofuginone to target stroma in a microfluidic device modeling the TME in pancreatic ductal adenocarcinoma increasing immune infiltration that was impaired by pancreatic stromal cells [62]. Halofuginone plays an antitumor role by inducing tumor apoptosis and autophagy, inhibiting cancer cell proliferation and metastasis, and cell cycle arrest through different signaling pathways such as TGF-β, caspase inhibition, and collagen synthesis inhibition [61].

In terms of acellular components of the TME and its spatial organization, we have validated our model to recreate the spatial arrangement and characteristics of the OC-TME and its influence on drug resistance. In this study, we have demonstrated that the incorporation of cancer-associated fibroblasts can modify cancer cell behavior, progression, invasion, and drug response. First, the endothelial cells grown in our model in the presence of VEGF were able to generate vessel-like structures mimicking the imperfect leaky vasculature in the TME [23, 24, 53]. In the presence of CAF, the inner diameter and elongation of the vessels were increased confirming that the CAF enhance the leaky vascularization and angiogenesis in OC-TME [10, 63]. Also, we confirmed that CAF are key contributors to ECM remodeling by higher collagen I secretion compared to normal stroma (NF and EC) and therefore converting the ECM in a physical obstacle with higher stiffness that impairs the penetration of drugs and oxygen into the tumor. We were able to recreate hypoxic tumor core characteristic for OC to mimic the role of hypoxia in drug resistance [64] and we validated that the presence of CAF in the OC stroma generated spatial gradients of oxygen from circulation chambers towards the cancer chamber compared to other stromal cells sole recreation of core hypoxic niches in the central chamber. Moreover, it is known that CAF secrete metabolites to fuel cancer cell growth under hypoxic and undernourished conditions and therefore increase tumor progression and invasion [37]. In our studies, we compared KURAMOCHI cells known for their low migratory potential and generation of clusters, and the SKOV-3 cell line with high metastatic potential [33]. As expected, a higher formation of the number of clusters in KURAMOCHI and higher migration in SKOV-3 were observed in the absence of stroma as there is no physical barrier caused by stromal cells, therefore lower gradients, and competition for nutrients with other cell types. It should be noted that is an experimental culture control condition, but not physiologically relevant. In the presence of stroma, CAF induced a higher formation of clusters than NF confirming their pro-tumorigenic potential [37]. In the migration studies of SKOV-3 cells, EC induced higher migration than when both types of fibroblasts were present, and of both stromal cocultures, migration in CAF was higher than NF. These effects are expected as it is known that both CAF and especially endothelial cells influence cancer invasiveness and metastasis in OC [9, 10, 23, 65]. In these studies, we observed that the cancer cells could interact with the stromal cell through the gaps between the pillars that separates the different compartments and recreate the tumor-stroma interactions, and also that the NF cultured for 14 days had low viability meanwhile CAF could still proliferate after 14 days in the tumor-on-a-chip thus the rest of the studies were performed for a maximum of 7 days in order to guarantee the viability of all cell types. Moreover, our microfluidic device allowed us to generate drug gradients as there was a delay of drug penetration in the cancer chamber compared to the stromal chamber, recapitulating the drug gradients in the TME, and allowing a laminar flow of the liquid injected in the circulation channel confirming the ability to study drug pharmacokinetics in our microdevice. Moreover, we confirmed that all cell types seeded in our microfluidic device could uptake the drug after 24 h of incubation, indicating our device is a suitable platform for drug screening that could be used in preclinical studies in a time-optimal manner. Moreover, we have validated that our model could mimic the tumor-stroma interactions and the influence of stroma in drug response to standard-of-care therapy (combination of paclitaxel and carboplatin) in OC [66]. As expected, normal stroma did not significantly influence the drug response in OC meanwhile the presence of CAF in the stromal compartment had a negative effect and caused drug resistance when compared to normal fibroblasts validating that the tumor-on-a-chip mimics the drug resistance mechanisms produced by CAF. Finally, we have demonstrated that targeting TGF-β signaling with halofuginone can rescue the drug resistance to paclitaxel and carboplatin caused by CAF in KURAMOCHI cells. Critically, these findings suggest that tumor-stroma interactions with CAF and endothelial cells and the biophysical properties of the OC-TME including hypoxia, ECM remodeling, and drug penetration are key contributors to OC chemoresistance. Moreover, we demonstrated that a precise recreation of the cellular crosstalk in the TME, as well as spatial organization through compartmentalization, are fundamental to determining the effect of these physical and biological mechanisms on drug resistance.

Currently, many studies have been conducted to assess the influence of the stroma in drug resistance in OC using different 3D preclinical models including spheroids, tumoroids, and 3D coculture models [53]. However, they do not recreate the compartmentalization, and dynamic properties of the TME [23, 26]. Therefore, microfluidic devices could bridge the gap between the preclinical *in vitro* and *in vivo* models as they are easy to use, high throughput, cost effective, and biocompatible, and allow to recapitulate compartmentalized TME, cell-cell interaction, gradients, cellular characteristics, migration, and fluid flow control [19, 27, 67, 68]. Several groups had developed microfluidic devices to assess the OC poor prognosis and study tumor-stroma interactions and migration of OC cells [69], biomarkers for early diagnosis [70], and shear stress [71]. For example, Ibrahim *et al*. developed a microchip to study the role of different stromal cell types such are mesothelial cells, endothelial cells, and adipocytes to model metastatic OC into peritoneum, but they did not model the primary OC tumors [46]. Dadgar *et al.* used OC patient-derived xenograft (PDX) tumors to generate spheroids in a multichambered microfluidic device demonstrating that the cell viability and epithelial markers in the cells grown in the microfluidic device were higher than in Matrigel 3D cultures but the generation of PDX models can be complicated, time-consuming and expensive and also it doesn’t meet the 3R (replacement, reduction, and refinement) for animal research [18, 72]. Alkmin *et al*. studied drug response to carboplatin in spheroids of different OC cell lines in channel-based microfluidic design with different collagen fiber morphology but they did not consider any stromal cell component [73]. Even though a similar approach was used to study tumor-stroma interactions in the microfluidic models for other cancer types such are breast cancer [74-76], pancreatic cancer [62, 77], glioma [45], colorectal [78], etc. to our knowledge, our model is the first microfluidic model for ovarian cancer that takes into account the spatial distribution of the key cell types present in its heterogeneous and complex TME, their interactions, and ECM remodeling and also recapitulates the hypoxic TME, allowing to study their influence in the drug resistance and targeting therapies that could improve the OC patient outcomes that have not significantly changed in the last 50 years [1].

Despite the fascinating findings in our study, there are some limitations that we would like to acknowledge. We used cell lines to model the OC-TME, the incorporation of primary cells would be more physiologically relevant and would recapitulate better individual drug response in OC patients allowing personalized therapeutic efficacy prediction [19]. We acknowledge that the hydrostatic pressure difference for laminar flow generation inside the microfluidic device lacks dynamic control and there is a pressure drop as the media flows through the channel. Alternatives to control fluid shear stress and fluid flow should be investigated [79]. Additional CAF- or ECM-targeting drugs or combination of drugs [66] should be further explored, as well as validation of TME compartmentalization with additional cell types present in the OC-TME such as adipocytes, mesothelial cells [46], and immune cells. Investigating tumor-immune interactions and immunotherapies could improve outcomes for OC patients, considering that high grade serous carcinoma was one of the first cancer types were the presence of tumor infiltrated lymphocytes was correlated with higher survival rates and CAF act as a barrier for immune infiltration and induce tumor inflammatory processes [28, 37].

## Conclusions

In conclusion, our results present a functionally characterized microvascularized multiniche tumor-on-a-chip able to recapitulate key spatial OC-TME compartmentalization including vascularization, ECM remodeling, oxygen gradients, and drug penetration kinetics, which directly influence drug resistance. Using this device, we have confirmed CAF’s role in modulating drug response and implemented targeted therapeutic approaches to overcome this chemoresistance. Our results are expected to have an important positive impact because they will provide strong evidence for providing new opportunities for improving chemotherapy effectiveness and mitigating chemoresistance in ovarian cancer.

## Acknowledgements

This project used Sanford Research Histology and Imaging Core and the Flow Cytometry Core Facilities that are supported in part by a Center for Cancer Research CoBRE grant from the National Institutes of Health (5P20GM103548). We want to thank Jared Wollman and Kelly Graber from the Sanford Research Flow Cytometry and Histology and Imaging Cores, respectively for training and help in experimental set up. Research reported in this publication was supported by the National Cancer Institute of the National Institutes of Health under award number R21CA259158 (P.P) and Pilot and Feasibility Enabling Technologies Award from Sanford Research and Sanford Health Foundation (P.P). E.S was funded by SPUR summer program under award R25HD097633. The content is solely the responsibility of the authors and does not necessarily represent the official views of the funding institutions.

