## Supplementary data for "Multicompartmentalized microvascularized tumor-on-a-chip to study tumor-stroma interactions and drug resistance in ovarian cancer"

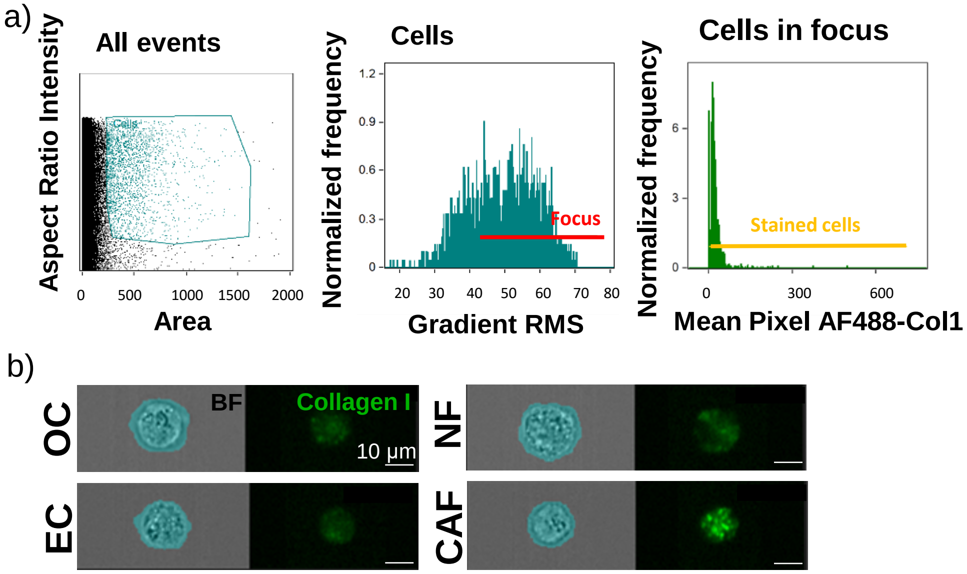


**Supplementary Fig. 1.** a) Gating strategy for imaging flow cytometry data using IDEAS v6.2 analysis software to detect from left to right the single round cells (brightfield area vs aspect ratio), in focus events (brightfield gradient RMS) and AF488-collagen I stained cells. b) cell mask (blue) used to quantify the MFI of AF488-collagen I inside the cells. Scale Bar=10 µm.

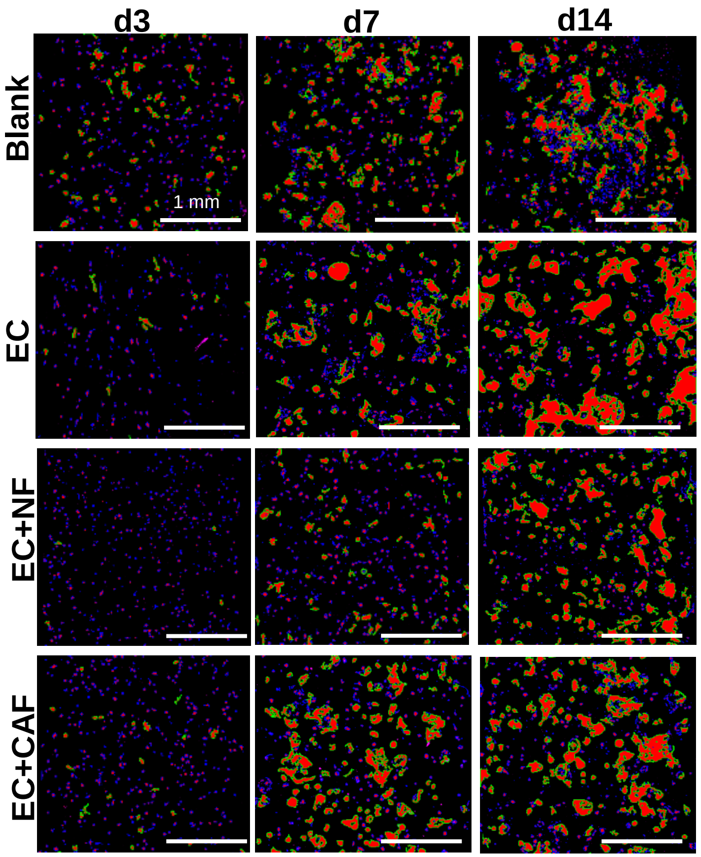


**Supplementary Fig. 2. Tumor-on-a-chip allows for cluster formation.** Representative close-up images of clusters of KURAMOCHI cells in cancer chamber on days 3,7 and 14, detected with NIS AR analysis software by binary thresholding and applying area restrictions in a range 50% higher than the calculated mean value of all objects and the largest object area (1300-90000 µm$^2$). Scale Bar= 1 mm.

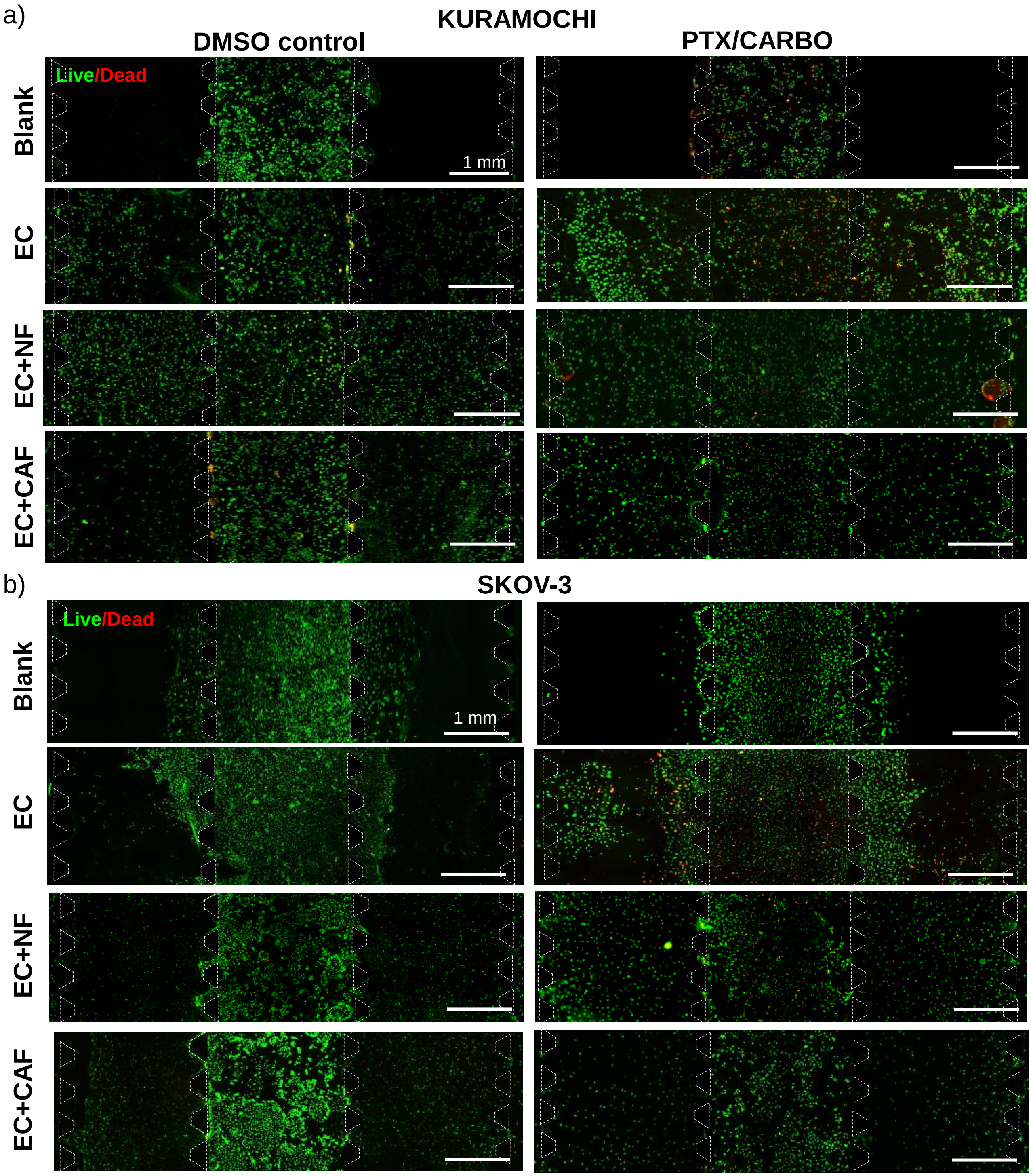


**Supplementary Fig. 3. Multiniche microvascular tumor-on-a chip recapitulates CAF-induced drug resistance in OC-TME. a**) Representative images of cancer and stromal chambers in Kuramochi or b) SKOV-3 (4M/ml) OC cell lines grown in central chamber with blank scaffold, EC, EC in co-culture with NF or CAF in stromal chamber for 3 days to allow ECM remodeling and then treated with PTX/CARBO combo (right) for 4 days. DMSO-treated microchips were used as a control (left). Live/Dead Nuclear ID dye was used to detect live cells (green) and dead cells (red). Scale Bar= 1 mm.

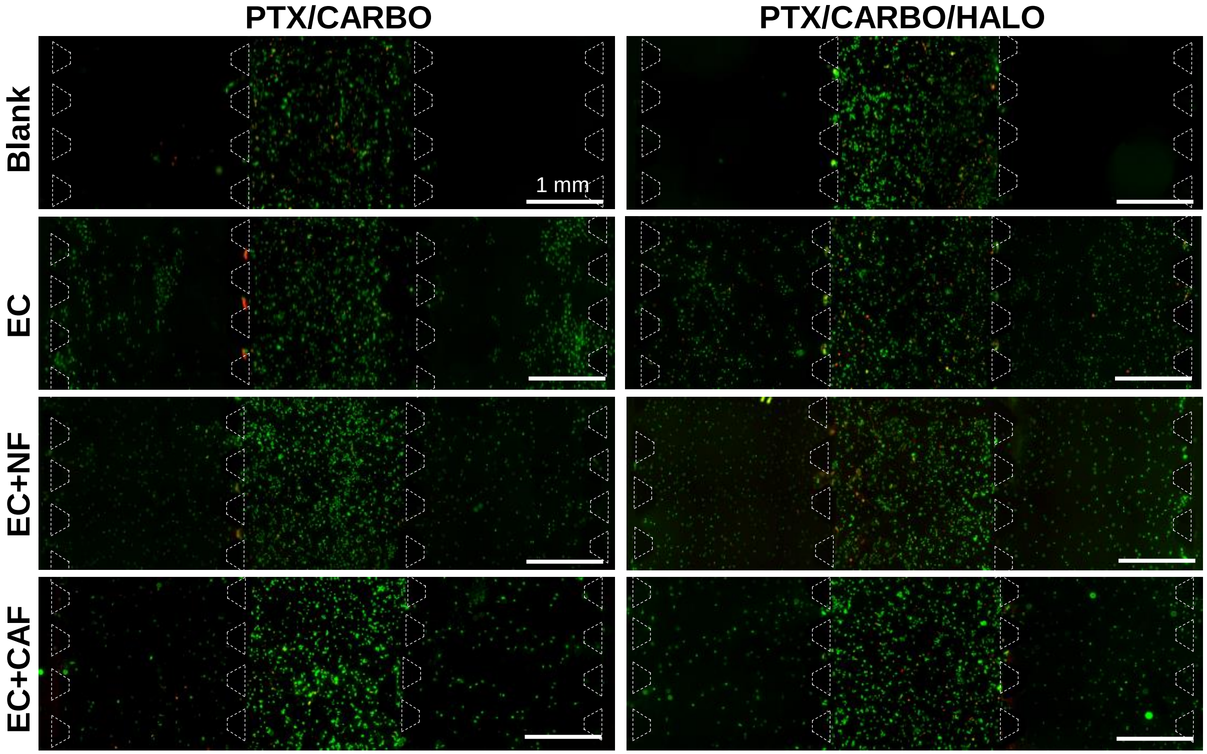


**Supplementary Fig 4. Drug resistance can be rescued by an anti-TGFb targeting agent halofuginone.** Representative images of cancer and stromal chambers of Kuramochi OC cells grown in central chamber with blank scaffold, EC, EC in co-culture with NF or CAF in stromal chamber treated with PTX/CARBO (left) or PTX/CARBO/HALO (right). Live/Dead Nuclear ID dye was used to detect live cells (green) and dead cells (red). Scale Bar= 1 mm.
